# Collective bacterial condensation is fundamentally constrained by the emergence of active turbulence

**DOI:** 10.1101/2025.07.20.665738

**Authors:** Nir Livne, Ady Vaknin

## Abstract

Collective bacterial condensation arises from positive feedback between the ability of bacteria to generate chemical gradients in their environment and their chemotactic ability to follow those gradients. This feedback drives the spontaneous formation of local cell accumulations, characterized by sharp cell-density gradients, even in the absence of physical boundaries. By following the dynamics of bacterial condensation in uniform acidic environments, we show that condensation is critically constrained by the spontaneous emergence of correlated bacterial swimming and the associated active turbulence. These collective behaviors generate vortex-like cell motion with a pronounced radial component directed down the cell-density gradient. This, in turn, induces local fluid motion that broadens the condensate and expels non-chemotactic bacteria. When condensates are strongly confined in thin layers, the radial component of fluid motion diminishes and condensation is enhanced. Moreover, in porous environments, correlated bacterial motion is strongly suppressed, allowing spontaneous condensation to progress even further until it reaches the limits imposed by the non-local nature of bacterial chemotaxis. Overall, these findings highlight the fundamental interplay between self-generated bacterial condensation and correlated swimming.

**Significance statement:** Self-induced condensation and active turbulence are two prominent forms of collective bacterial behaviors. However, their interplay has not been experimentally studied. Here, we show that collective bacterial condensation gives rise to correlated swimming, which in turn fundamentally limits further condensation. Furthermore, we demonstrate that in dilute porous environments—common in natural bacterial habitats—correlated swimming is strongly suppressed, enabling the spontaneous formation of extremely dense cell condensates. These condensates may serve as a basis for the development of more structured bacterial communities.

## Introduction

Motile bacteria navigate their environment by tracking chemical gradients toward regions they deem favorable (1–4). This behavior, termed chemotaxis, influences many aspects of bacterial ecology (5, 6) and human health (7–9). Even in spatially uniform environments, bacteria can collectively follow self-generated gradients, enabling them to accelerate their expansion into resource-rich regions (10, 11), or alternatively, to condense into focal points by secreting beneficial (attractant) effectors (12, 13) or neutralizing harmful (repellent) effectors (14).

Notably, in contrast to the inherently dynamic bacterial accumulation triggered by exogenous chemoeffector release (15) or observed in expanding bacterial bands (16), bacterial condensation can lead to sharp but quasi-static accumulations (14). This condensation relies on positive feedback between self-generated gradients and the chemotactic responses of the bacteria to those gradients (Fig. 1A), which, in principle, can lead to a divergence in the local cell density (17). Studies of transient and low-density cell accumulations limited by oxygen depletion have suggested that a fundamental limit on such cell condensation arises from the fact that bacterial gradient sensing itself is a non-local behavior (18). However, the constraints on bacterial condensations under conditions of oxygen abundance remain unclear. Here, we show that under these conditions, bacterial accumulations indeed progress further, forming stable condensates that are nevertheless limited by the emergence of correlated bacterial swimming and the associated active turbulence.

**Figure 1.**
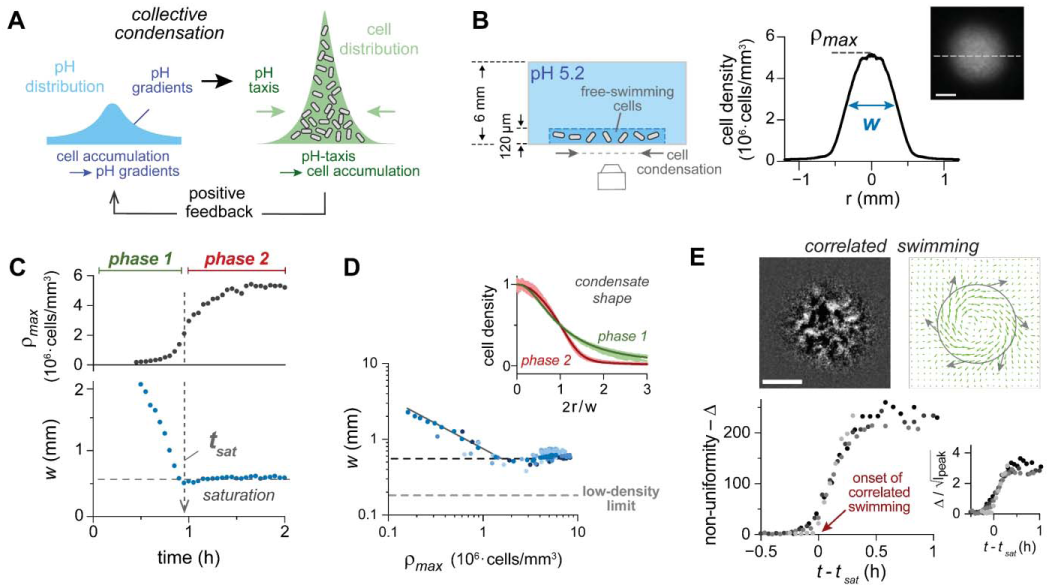
Bacterial condensation is constrained by correlated swimming. (A) Schematic of bacterial condensation in acidic environments. Bacteria locally neutralize the medium and generate pH gradients within an otherwise uniform acid medium. At the same time, cells are chemotactically attracted toward neutral pH. Together, these behaviors form reinforcing feedback that drives bacterial condensation. (**B**) The experimental setup consists of a thin suspension of GFP-expressing cells (5 mm diameter) confined in a circular well formed in 3% agarose gel. Both the cell suspension and the surrounding gel initially form a uniform acidic environment (pH 5.2). Condensation was monitored using fluorescence microscopy. Shown on the right is a typical mature condensate 2 h after initiation (scale bar, 300 μm), along with the radially averaged cell-density profile. The peak cell density (*ρmax*) and the width at half maximum (*w*) are indicated. (**C**) A typical condensation dynamic, characterized by increasing *ρmax* and decreasing *w*, until *w* abruptly saturates at time *tsat*. (**D**) The condensate width (*w*) is plotted against peak cell density (*ρmax*) for seven independent experiments. The solid gray line represents a power-law slope of -2/3. The saturation value of *w* (dark-gray dashed line) and the corresponding value measured at low cell density (light-gray dashed line; see ref. 18) are indicated. *Inset* – normalized cell density profiles from three independent experiments; profiles taken at *t < tsat* (Phase 1) are shown in green, and profiles taken at *t > tsat* (Phase 2) are shown in red. Each profile was normalized by subtracting the background intensity and dividing by the peak intensity; the radial coordinate was normalized by *w*/2. (**E**) *Upper-panel* – a representative (filtered) fluorescence image of the condensate, demonstrating the dynamic non-uniformity developed at high cell density (left, see Materials and Methods and Movie S1; scale bar, 300 μm), along with the corresponding time-averaged (30 s) velocity field obtained by particle image velocimetry (right, gray arrows are a guide to the eye). *Lower panel* – the condensate non-uniformity (Δ), evaluated within the region r ≤ 300 μm, is plotted over time for four independent experiments (see Materials and Methods). *Inset* – The relative non-uniformity (Δ /jl_peak_) plotted over time.

*Escherichia coli* cells swim by rotating helical filaments (flagella) that bundle and push the cell forward, whereas during stochastic episodes of counter-rotation, the bundle comes apart, causing random reorientation. These two swimming modes—forward swimming (‘run’) and random reorientation (‘tumble’)—together generate an active random walk with a typical run length of ∼10–30 μm (19). In the presence of effector gradients, the probability of tumbling is modulated to bias the random walk toward beneficial effectors and away from harmful ones. In particular, bacteria can migrate toward neutral pH (20, 21), and, combined with their ability to locally neutralize the acidic environment, this behavior leads to spontaneous condensation (14). Such condensation is readily observed when the bacterial suspension is coupled to a large, cell-free environment that maintains the pH level within the dynamic sensory range and prevents oxygen depletion, which is required for motility. The number of condensates depends on the lateral size of the bacterial-suspension layer: from quasi-ordered arrays of bacterial condensates in large suspensions spread on agar plates to single condensates in layers only a few millimeters in diameter (14).

On the other hand, at high cell densities, bacteria tend to exhibit correlated swimming and flocking-like behavior (22–24), which can lead to turbulent-like motion often termed ‘active turbulence’ (25, 26). Such dynamics arise from direct steric and hydrodynamic interactions between cells. With notable exceptions (27, 28), correlated bacterial swimming has mostly been studied in uniformly distributed populations confined by physical boundaries. In contrast, bacterial condensation leads to highly non-uniform cell distributions in the absence of explicit physical confinement. Numerical studies have suggested that dilute self-attracting run-and-tumble particles with weak chemotactic strength aggregate less efficiently when self-induced fluid motion is considered (29). Experiments have indicated that cell alignment may constrain the angular reorientation of individual cells and reduce chemotactic efficiency in uniform cell suspensions and shallow effector gradients (30). Other studies, however, have shown that the tendency of cells to align may, in fact, enhance their taxis efficiency toward a local source (31). Overall, the interplay between bacterial condensation and correlated swimming has been scarcely explored, and how correlated bacterial swimming influences their collective condensation remains unclear.

Here, we investigate the limits of bacterial condensation at high cell densities under conditions where oxygen deprivation and effector saturation are not limiting factors. As a model, we used a recently described assay for bacterial condensation in uniform acidic environments (14), under conditions that allow us to follow the dynamic development of individual condensates. We show that bacterial condensation is sharply constrained by the emergence of correlated bacterial swimming during cell accumulation. This correlated swimming gives rise to substantial turbulent-like fluid motion, characterized by vortex-like motion with an additional outward velocity component directed down the cell-density gradient. Such fluid motion exerts outward physical forces on the cells that counteract the inward chemotaxis force, thereby broadening the condensate and expelling non-chemotactic cells. A simple model incorporating cell-induced radial outflows quantitatively reproduces these observations. Moreover, suppressing this outward fluid motion in very thin bacterial layers leads to enhanced condensation, whereas in porous environments, correlated bacterial motion is suppressed altogether and condensation is further enhanced, approaching the low cell-density limit but with significantly higher cell densities.

## Results

### Bacterial condensation saturates at the onset of correlated swimming

To test the constraints on high-density bacterial condensation in a controlled environment, we used the setup illustrated in Fig. 1B (14). In this setup, free-swimming cells are confined to a shallow liquid layer (typically ∼120 µm high) between a glass coverslip and a thick hydrogel layer (3% agarose; 6-mm high), such that, initially, the bacterial suspension and the hydrogel together form a uniformly acidic environment (pH 5.2). While cells cannot penetrate the dense hydrogel mesh, protons and oxygen readily exchange between the thin bacterial suspension and the gel, preventing their rapid depletion inside the bacterial condensate and thereby enabling quasi-stable, high-density cell condensation (Fig. 1B, right plot). To study the dynamics of individual condensates, the lateral size of the bacterial layer was restricted to 3-5 mm.

By following the peak cell density (𝜌_𝑚𝑎𝑥_) and the width (*W*) of the condensate, we identified two distinct temporal phases of condensation (Fig. 1C). Initially, in phase 1, the peak cell density 𝜌_𝑚𝑎𝑥_ increased while the width *W* decreased, indicating strong bacterial condensation. Then, beyond a critical time point (t_sat_), in phase 2, the width *W* abruptly plateaued while 𝜌_𝑚𝑎𝑥_ continued to rise, reflecting continued cell accumulation without further narrowing of the condensate. These dynamics are further demonstrated in Fig. 1D, where the condensate width is plotted against the peak cell density. This plot reveals a power-law dependence 𝑤∼𝜌_max_^−2/3^ in phase 1, followed by a clear saturation of the condensate width at a characteristic cell density. Interestingly, the saturated width is markedly larger value than the low-cell-density limit previously reported (18), where condensate width corresponded to only several run lengths. The spatial profiles of the condensates also reflected the dynamic transition between phases 1 and 2 (Fig. 1D, inset). When each profile was normalized by 𝜌_𝑚𝑎𝑥_ and the radial coordinate scaled by the width *W*, the profiles collapsed onto a single curve within each phase but were clearly distinct between the two phases.

The dynamic transition between phases 1 and 2 was clearly associated with the appearance of significant density fluctuations within the condensate (Fig. 1E). Initially, in phase 1, the bacterial condensate was smooth and stable (Fig. S1). However, beyond the critical time point t_sat_, in phase 2, a pronounced spatial non-uniformity developed on top of the average cell-density profile, with dynamic fluctuations emerging at spatial scales of ∼ 20-100 μm (Fig. 1E and Movie S1; see Materials and Methods), substantially larger than individual bacteria. These cell-density fluctuations indicate correlated swimming of large groups of cells that, on average, exhibit collective motion characterized by a vortex-like rotation of the entire condensate together with an additional outward-directed radial component (Fig. 1E; see Materials and Methods).

Notably, in all cases, the vortex rotation was consistently clockwise, as viewed from the glass side.

### Fluid motion associated with correlated bacterial swimming

To characterize fluid motion generated by correlated bacterial swimming in phase 2, we mixed green fluorescent beads (1 μm in diameter) with the bacterial population, which in this case expressed red fluorescent proteins, and analyzed bead motion by particle tracking (Materials and Methods). Typically, the bead trajectories consisted of periods of local motion interrupted by approximately directional movements, termed ‘jets’(25) (Fig. 2A). The influence of the bacterial motion on the beads was evaluated for each jet by the ratio *Pe* = v·L/D_beads_ (Péclet number), where v and L are the velocity and length of each jet, respectively, and D_beads_ is the diffusion coefficient of the beads. This ratio compares advective and diffusive transport; v/L and D_beads_ /L^2^, respectively. The cumulative distribution of *Pe* is shown in Fig. 2A (right plot) for jets located within (green) or outside (blue) the condensate region. Clearly, bead motion was enhanced within the condensate and was dominated by advection (Pe≫1).

**Figure 2.**
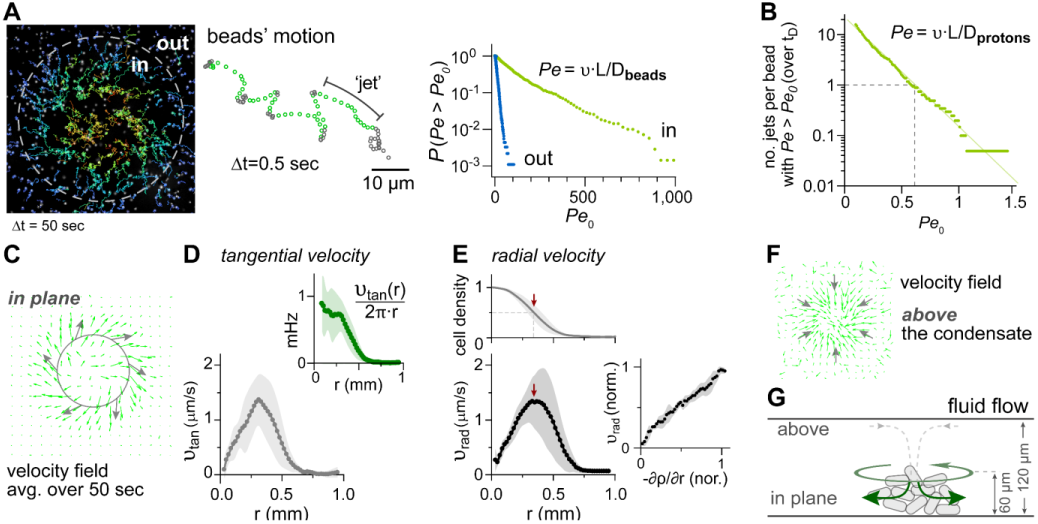
Fluid motion within mature condensates. Fluid motion was assessed by tracking 1 μm fluorescent beads (green) mixed within the bacterial suspension. In this case, cells were labeled with mCherry (red). (**A**) Traces of individual beads (over 50 sec) within a mature condensate, captured using particle-tracking analysis (Materials and Methods). The dashed white circle indicates the condensate width (*w* ∼ 550 μm). A representative trace is highlighted on the right, illustrating intermittent motion composed of periods of local random motion (gray symbols) and periods of directed motion, termed ‘jets’ (green symbols). The length (*L*) and average speed (*v*) of each jet were used to calculate *Pe = v·L/Dbeads* (with D*beads* = 5·10^-9^ cm^2^ s^-1^). The cumulative distribution of *Pe* is shown on the right for beads inside or outside the contour defined by the condensate width. (**B**) Comparison of bead motion with the expected proton diffusion. First, the number of jets per bead with a *Pe* value larger than a given threshold (*Pe0*) were found and then multiplied by the typical diffusion time of protons across the condensate tD ∼ w^2^/4DH+ ∼ 4min (using DH+ = 5·10^-6^ cm^2^ s^-1^). (**C**) Time-averaged velocity field in the condensate plane, extracted using PIV (Materials and Methods). (**D**) Time-averaged tangential motion inside condensates (mean and standard deviation over five experiments). *Inset* – angular velocity; positive values correspond to clockwise rotation. (**E**) Radial velocity (bottom) and the corresponding normalized cell density profile (top; mean and standard deviation over five experiments). The red arrow marks the condensate width (w). *Inset* – radial velocity plotted against the local cell density gradient. (**F**) Measured velocity field above the condensate (h ∼100 µm), showing an inward radial flow. (**G**) Schematic summarizing the inferred fluid motion in the vicinity of the condensate (see also Movie S2).

To further evaluate the potential significance of bacteria-induced advection on effector mixing, we computed the proton Péclet number, *Pe* = v·L/D_H+_, assuming that protons follow the local fluid motion. Because proton have a much larger diffusion coefficient than the beads (D_H+_ ∼ 1000·D_beads_), advection of protons is expected to be less effective. Nevertheless, we find that during the time it takes for a proton to diffuse across the condensate t*_D_* = W^2^/(4*D_H+_*), each tracer bead in the condensate experiences, on average, at least one substantial jet with *Pe* > 0.5 (Fig. 2B). Thus, cell-induced advection can have only a moderate influence on proton distribution.

Time-averaged bead motion within the condensate also exhibited a clockwise vortex-like motion with a radially outward component (Fig. 2C–E; analyzed by particle image velocimetry, PIV, see Materials and Methods). The tangential velocity depended on the distance from the center of the condensate, such that the condensed bacteria exhibited a vortex-like rotation at ∼1 mHz (Fig. 2D). The outward radial component extended throughout the condensate and scaled approximately linearly with the local cell-density gradient (Fig. 2E). Notably, the peak radial velocity (∼ 1.5 μm·s^-1^) is comparable to the chemotactic drift velocity of these cells (19, 30). Finally, the condensed cells tended to occupy the lower portion of the fluid layer, and above the condensate the tracer beads exhibited an inward motion (Fig. 2F; Movie S2). Thus, the fluid motion near the condensate resembles a three-dimensional toroidal pattern (Fig. 2G).

### Effect of bacterial-layer height and porous environment

To further examine the interplay between bacterial condensation and correlated swimming, we sought to physically constrain correlated swimming (Fig. 3). This was done in two ways: first, by confining the bacteria to thinner layers whose height is less than the condensate height (*h_0_* ∼60 μm) that spontaneously formed in *h* ∼120 μm layers (Fig. 2G); and second, by embedding the bacteria in a porous environment, which is expected to dampen fluid flows within the condensate (32) and has been suggested to inhibit bioconvection (33).

**Figure 3.**
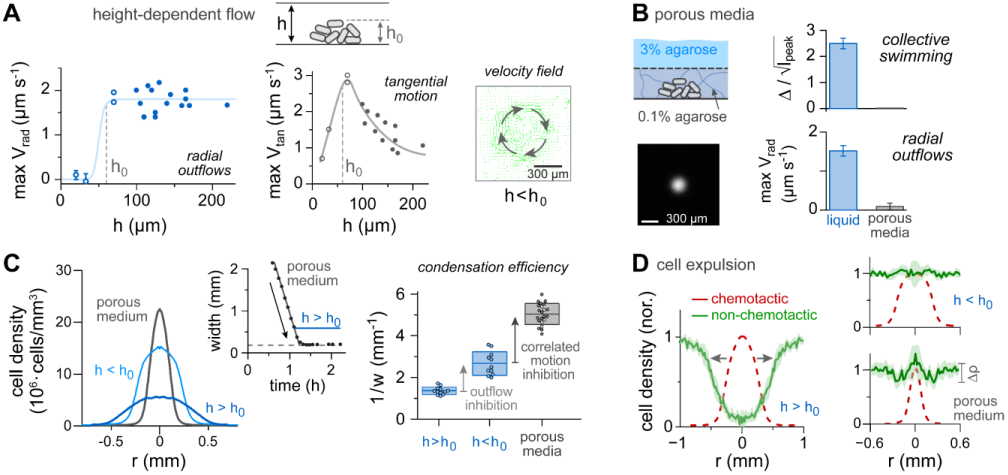
Inhibition of collective bacterial motion enhances condensation. (**A**) Height dependence of the bacterial-induced fluid motion. Condensation experiments in the presence of fluorescent beads performed using varying heights of the bacterial layer (h) and initial (uniform) cell density of 0.4·10^6^ cells/mm^3^. Time-lapse images of beads were taken using 10x (open symbols) and 4x (filled symbols) objectives and analyzed using PIV. The maximal radial (left) and tangential (right) velocities are shown for mature (stable) condensates. The condensate height observed in experiments with h ∼ 120 µm is defined as h0 (∼ 60 µm; see Fig. 2). (**B**) Bacterial condensation in porous media. Condensation experiments were conducted in a device similar to that shown in Fig. 1B, except that the bacterial suspension was embedded within a low-concentration (0.08*–*0.12% w/v) agarose hydrogel. A schematic illustration of the device, along with a representative fluorescence image of a mature condensate are shown on the left. The right panel compares collective bacterial swimming and the radial outflow observed in the presence or absence of the agarose hydrogel (mean ± SD, N=3). Collective swimming was quantified by the relative non-uniformity of the condensate (see Fig. 1). (**C**) *Left plot –* Representative cell-density profiles in mature condensates obtained in liquid environment with h < h0 or h > h0 and in a porous medium with h > h0 (as labeled). *Right plot –* quantitative comparison of condensation efficiency— defined as the inverse width (1/w)—in the various environment (box plots show mean and SD for N = 12, 11, 22 (left to right). In porous media, filled and open symbols represent data in 0.08% and 0.12% agarose (w/v). *Inset (center) –* condensation dynamics in porous medium (black symbols), shown together with the saturation value in liquid (blue line; h > h0) and the corresponding value observed in ref. 18 for experiments with very low cell-density (gray dashed line). (**D**) Expulsion of non-chemotactic cells from condensates. Condensation experiments of mCherry-labeled chemotactic cells were repeated in the different environments (as labeled) while adding GFP-labeled non-chemotactic cells (ΔcheY/Z) to the initial (uniform) cell suspension, at a 100:1 ratio, respectively. The cell-density profiles of the non-chemotactic cells in a mature condensate are shown in green (solid line ± SD, N = 3) along with the corresponding profile of the condensate (red dashed line). Profiles of non-chemotactic cells were normalized to the density far from the condensate, and profiles of the chemotactic cells were normalized to the peak density. Δρ (lower-right plot) represents the SD of variability in the distribution of non-chemotactic cells within each experiment.

The effect of bacterial-layer height on the time-averaged flow velocities within mature condensates at saturation is shown in Fig. 3A. Interestingly, the maximal radial outflow diminished sharply for heights smaller than h_0_, while the collective motion persisted (Movie S3), resulting in dominant tangential rotation for h < h_0_. This effect may arise from the elimination of the space above the condensate where the opposing radial fluid inflows were observed (Fig. 2G). The tangential velocity was also modulated by the height of the bacterial suspension, peaking at approximately h ∼ h_0_, though the origin of this behavior remains unclear.

The effect of porous environments on correlated bacterial motion within mature condensates at saturation is shown in Fig. 3B. Experiments were conducted using a setup similar to that shown in Fig. 1, except that the bacteria were immersed in a porous agarose hydrogel environment (0.08–0.12% w/v). These agarose concentrations were chosen, on the one hand, to minimize perturbations to motility and chemotaxis and, on the other hand, to still allow a structured mesh (34). The chemotactic performance and the effective diffusion of bacteria within such hydrogel layers were similar to those observed in liquid (Fig. S2), and clearly supported cell condensation with even similar dynamics in phase 1 (Fig. 3B–C).

However, porous environments suppressed the collective swimming and the associated fluid flows (Fig. 3B).

The corresponding influences of the suspension height (h) and porous environments on bacterial condensation are shown in Fig. 3C. Clearly, bacterial condensation becomes progressively narrower as correlated swimming is constrained: first, by reducing the suspension height, which mainly diminishes the radial motion; and second, by introducing a porous environment, which eliminates correlated swimming altogether. Notably, condensation proceeded in porous media at a rate similar to that in liquid (middle plot), but it saturated only when it approached the low-density limit (∼180 μm), associated with the non-local nature of chemotaxis (18). These condensates were also more compact in the vertical direction, spanning only ∼20–30 μm (Fig. S3), and thus their local cell density is approximately five times the values indicated in the plots, which is averaged over the bacterial layer. Thus, alleviating the constraints related to oxygen and collective bacterial motion allowed condensation to reach its basic limit imposed by the inherent non-locality of chemotaxis (18); however, here the condensing bacteria reach this limit at much larger cell density, corresponding to a volume fraction of approximately 0.25 (Fig. S3), while the cells remained largely motile (Movie S4).

### Condensation-induced cell expulsion

The correlations observed between collective motion and condensate broadening (Fig. 3A–C) suggest that correlated motion may exert an outward force on the condensing bacteria. To examine this possibility, we repeated the condensation experiments while mixing the chemotactic cells (red-labeled) with non-chemotactic (*cheYcheZ)* cells (smooth-swimming; green-labeled), which are not subjected to the inward chemotactic force and therefore serve as a probe (Fig. 3D). Indeed, in liquid suspensions with h > h_0_, which exhibit the weakest condensation, the non-chemotactic cells were clearly expelled from the condensate (Fig. 3D, left). Run-and-tumble but non-chemotactic cells were also tested (35) and similarly showed significant depletion (Fig. S4). In contrast, in liquid suspensions with h < h_0_, in which the radial outflows are suppressed, no substantial expulsion of the non-chemotactic cells was observed (Fig. 3D, right). Similarly, despite the strong condensation of the chemotactic cells in porous (hydrogel) environments, expulsion of the non-chemotactic cells was not observed in this case (Fig. 3D, right).

Taken together, these results suggest that expulsion of the non-chemotactic cells from the condensate in thick liquid suspensions is likely due to the radial outflows induced by correlated swimming, rather than merely to the locally elevated density of the chemotactic cells. Moreover, since similar outward forces are expected to act on the chemotactic (condensing) cells as well, these flows likely contribute to broadening the condensate itself.

### Ring-like condensates

Tight coupling of the thin bacterial suspensions to the surrounding cell-free hydrogel reservoir (Fig. 1B), together with low initial cell density used in the experiments described thus far (Figs. 1–3), prevents the condensates from reaching cell densities high enough to trigger substantial oxygen-dependent repellent responses and allow stable condensation. Replacing the glass coverslip in these experiments with an oxygen-permeable surface did not significantly affect condensation, further supporting the conclusion that oxygen limitation was not substantial (Fig. S5). However, when the initial uniform cell density was systematically increased, we observed that once sufficient cells accumulated in the condensate, it tended to adopt a ring- shaped structure that, in turn, limited the maximal cell density (Fig. 4A). Ring formation most likely resulted from local oxygen depletion by the condensing bacteria, which induces an outward aerotactic response (36–38). This behavior occurred in both liquid and porous environments.

**Figure 4.**
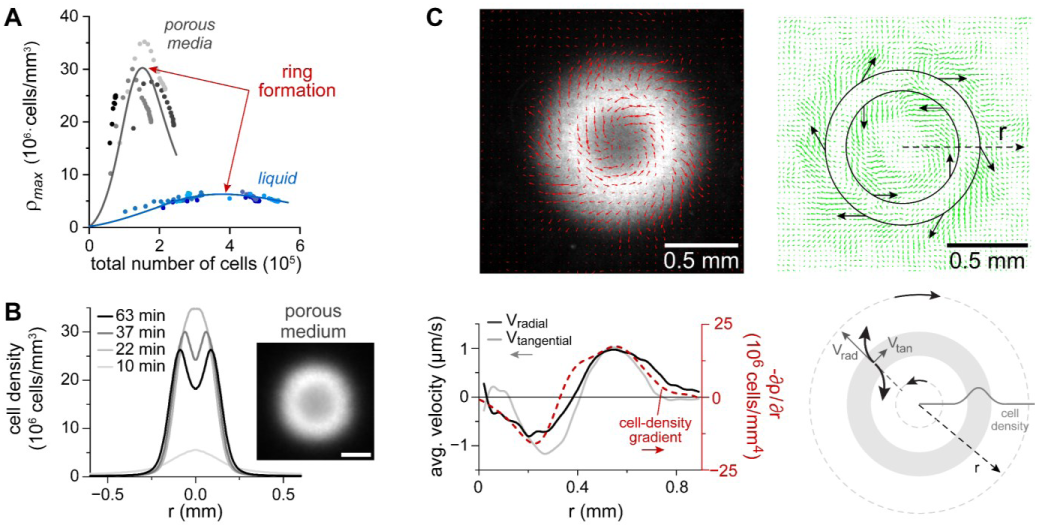
**Ring-shaped bacterial condensation**. Condensation experiments were performed for progressively higher initial (uniform) cell densities. (**A**) Peak cell density plotted against the total number of cells in the condensate, for experiments performed in liquid (h > h0; blue symbols) or in porous media (gray symbols). The decrease in peak cell density at high total cell numbers correlates with the formation of ring-shaped condensates. (**B**) Representative evolution of a ring-shaped condensate observed in porous environments. The condensate profile is shown (left), together with a fluorescence image taken at 50 min (scale bar, 150 µm). (**C**) Correlated swimming in ring-shaped condensates. A typical fluorescence image of a ring-shaped condensate in liquid medium (h > h0; upper-left image) and the corresponding time-averaged bacterial velocity field extracted using PIV (overlaid red arrows). For clarity, the same velocity field is shown separately on the right (green arrows). Below the image, the radial (light gray) and tangential (dark gray) velocities are plotted as a function of distance from the condensate center (left ordinate), together with the corresponding cell-density gradient (red dashed line; right ordinate). The correlated reversal of both velocity components across the ring is illustrated schematically (lower right), producing a consistent rightward swimming tendency in both inner and outer regions.

The ring-shaped condensate allowed us to uniquely examine the relationship between the correlated bacterial motion and the local cell-density gradient under conditions where the density is non-monotonic along the radial direction and its gradient changes sign. Interestingly, we found that both components of the correlated-motion vorticity reversed direction between the inner and outer regions of the ring (Fig. 4C). Thus, the radial motion aligns with the cell-density gradient but also exhibits a rightward lateral deflection relative to this direction (see illustration in Fig. 4C).

### Simplified model for condensate broadening

To test the minimal model capable of capturing the bacterial condensation dynamics observed here, we used a previously suggested Keller–Segel-based model for bacterial condensation in acidic environment (Eqs. 1–2, ref. (14, 39, 40)), augmented with a radial outward bacterial drift (Eq. 3) that arises from the correlated bacterial motion and is proportional to the local cell-density gradient, as shown in Figs. 2D and 4C.

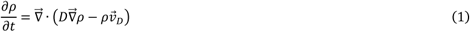

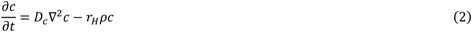

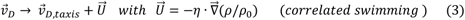

Here, 𝐷 is the effective diffusion coefficient of swimming cells, 𝐷_𝑐_ is the proton diffusion coefficient, 𝑟_𝐻_ = 𝛾𝑐^1/2^ is the proton-neutralization rate measured in ref. (14). 𝑣_𝐷_ is the bacterial drift velocity: in the absence of correlated swimming, 𝑣_𝐷_ only reflects chemotaxis, and in the presence of correlated swimming 𝑣_𝐷_ also includes 𝑈⃗, the effective in-plane drift bias imposed on the cells by the 3D fluid outflow (see Fig. 2G). The chemotaxis drift velocity was modeled as 𝑣 _𝐷,𝑡𝑎𝑥𝑖𝑠_ = 𝛽∇⃗ ln((𝑐 + 𝐾_𝑜𝑓𝑓_)⁄(𝑐 + 𝐾_𝑜𝑛_)) where 𝐾_𝑜𝑛_ and 𝐾_𝑜𝑓𝑓_ are the effective dissociation constants for the chemoreceptors in the ON or OFF states, and 𝛽 is the chemotaxis coefficient (41). The parameter η was extracted from the data in Figs. 2D and S7 (1.7 ± 0.5 x 10^-7^ cm^2^s^-1^), and ρ_0_ is the initial (uniform) cell density (∼0.2·10^6^ cells/mm^3^). Because diffusion across the thin bacterial layer is rapid (∼20 s) relative to bacterial condensation time, the bacterial dynamics was approximated as effectively two dimensional, coupled to the three-dimensional proton field. For more details see SI Model.

The basic model, in the absence of correlated swimming (𝑈⃗ = 0), captures the positive feedback between the tendency of cells to neutralize their environment (𝑟_𝐻_) and their tendency to migrate down repellent gradients (𝑣 _𝐷_), thereby driving condensation. This model can also account for the dynamics and shape of the condensates in phase 1; however, it predicts condensates in phase 2 that are markedly distinct from those observed experimentally (Figs. 5A–B and S6). Incorporating the outward drift component (𝑈⃗, Eq. 3), strictly limits the condensation and accounts for the broadening and overall shape of condensates in phase 2 (Fig. 5A-B, black lines and SI Model). Adding this drift component is included at the beginning of the numerical solution transition between phases 1 and 2 is more gradual than the observed transition (Figs. 1C and 5B). However, accounting for the fact that correlated motion emerges only above a threshold cell density (Fig. 1) by adding this drift component above a threshold density, clearly capture also for the shape transition (Fig. 5B). Moreover, this model also quantitatively account for the expulsion of non-chemotactic cells from the condensate (Fig. 5C and SI Model). Finally, the model can also quantitively account for the power-law dependence of the condensate width on its peak density (Figs. 1D and S6).

**Figure 5.**
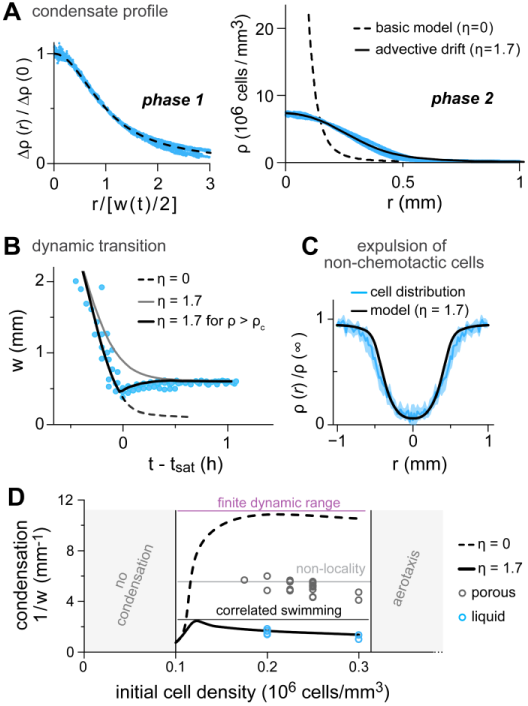
A model for condensate broadening by collective motion. Numerical solutions of the model described by Eqs. 1–3 were used to compare model predictions with experimental data obtained in liquid medium (h > h₀). The parameter η in Eq. 3 is expressed in units of 10⁻⁷ cm² s⁻¹. (**A**) Condensate profiles (blue symbols; N = 4, from Fig. 1D) measured in Phase 1 (left) or Phase 2 (right), shown together with model solutions for η = 0 (dashed line) and η = 1.7 (solid black line). (**B**) Time dependence of condensate width (blue symbols; N = 3, from Fig. 1), shown together with model solutions using constant η values of 0 or 1.7. To introduce correlated swimming only when ρmax > ρc, we set η(ρmax) = η0 / [1 + exp(−(k / ρ₀)(ρmax − ρc))], where ρ₀ is the initial uniform cell density, ρc = 25·ρ₀, and k = 3 (solid black line). (**C**) Data from Fig. 3D (liquid, h > h₀) are replotted (blue line) together with the distribution predicted by the model (black line): ρ = exp[−(η / DNC)(ρWT(r) / ρ₀)], with η = 1.7, ρWT(r) taken from the experimental data in (A), and DNC = 2.5 × 10⁻⁶ cm² s⁻¹ representing the effective diffusion coefficient of non-chemotactic cells (see derivation in SI Model). (**D**) Summary plot. The saturated condensation level (1/w) is plotted as a function of the initial uniform cell density for experiments in liquid (blue symbols; h > h₀) and in porous media (gray symbols), together with the corresponding model solutions for η = 0 and 1.7. At low cell densities, the uniform cell-density is stable (left shaded region; refs. 14, 40). At high cell densities, an aerotaxis-mediated ring appears that restricts condensation (right shaded region). Between these limits, condensation is first constrained by correlated bacterial swimming (black line), then by the non-local run-and-tumble mechanism of chemotaxis (‘low-density limit’, ref. 18), and ultimately by the finite dynamic range of bacterial sensory signaling (dashed black line; ref. 14).

The overall relationship between the initial cell density and condensation is summarized in Fig. 5D. When the initial density is below approximately 0.1·10^6^ cells/mm^3^, no condensation is observed (14, 40). At higher densities, the initially uniform distribution becomes unstable, and condensates emerge. In the absence of correlated swimming, bacteria would condense until the local cell density raises the pH beyond the sensory dynamic range of the cells (purple line, see also ref. (14)). However, because correlated swimming readily emerges, bacterial condensation is severely restricted (model: black line; data (h > h_0_): blue circles). In porous media condensation proceeds further (data: gray circles), though remains limited below the dynamic-range limit, likely due to the non-local nature of bacterial gradient sensing (18). At even higher initial cell densities, condensation becomes limited by ring formation, presumably due to aerotaxis (Fig. 4).

## Discussion

Chemotaxis-mediated condensation and correlated swimming are two prominent collective behaviors exhibited by motile bacteria. Here, we describe a fundamental interplay between these behaviors, whereby, bacterial condensation is strictly constrained by the emergence of correlated swimming at high cell density. Bacterial condensation proceeds in two phases (Fig. 1): initially, the population condenses and the local cell density increases; then, upon the onset of correlated swimming, the width of the condensate plateaus while cell density continues to rise, namely, bacteria accumulate but stop condensing. Bacterial condensates—characterized by steep cell-density gradients and a lack of physical boundaries— give rise to a distinct form of correlated swimming and active turbulence, with a radial component aligned down the cell-density gradient (Figs. 2 and 4). This radial component diminishes when the condensate is restricted by the height of the bacterial layer, and correlated swimming diminish altogether in porous environments, which ultimately allow for condensation to proceed and form a robust bacterial condensation with high cell density (Fig. 3). Similar interplay between bacterial correlated swimming and other forms of collective condensation can be generally expected.

With notable exceptions (28, 42), active turbulence has been mostly studied in uniform cell populations confined by well-defined lateral boundaries, either solid (43) or at liquid–air or liquid–oil interfaces (43–46). In such settings, bacterial behavior near the lateral physical boundaries may play an important role in shaping the observed swimming patterns (44, 47). In contrast, the bacterial condensate examined here represents a unique system with a self-sustained yet highly non-uniform cell distribution lacking lateral physical boundaries. The fluid motion in the vicinity of the condensate exhibits a three-dimensional toroidal pattern (Fig. 2F), reminiscent of the flow patterns observed in bioconvection (33, 48) or similar systems (49).

However, in bioconvection the flow arises from an instability caused by upward bacterial taxis opposing the downward gravitational force. In contrast, the bacterial condensate remains stably positioned near the lower part of the liquid layer, and only the fluid motion exhibits toroidal-like motion. Notably, disrupting this fluid motion in thin bacterial layers (h < h_0_) clearly affects the correlated motion.

By driving the expulsion of motile-but-non-chemotactic bacteria from the condensate, the emerging correlated bacterial motion effectively segregates the initially mixed population into chemotactic and non-chemotactic population (Fig. 3). We have previously shown that cell condensation can expels cell aggregates (10–20 µm in size), which are spontaneously form if cells are pre-grown at 37°C (14). Similarly, aerotaxis-driven accumulation of bacteria has also been shown to displace algal cells (50). Additionally, at higher cell densities, expulsion the bacterial cells themselves has been observed, which appeared non-motile (50). Other experiments have demonstrated bacterial expulsion from a vortex induced by a rotating magnetic microsphere at 2–40 Hz (27). Notably, the vortex generated here by cell condensates is much slower (∼10^-3^ Hz). Here, we find that the expulsion of motile-but-non-chemotactic cells from the condensate relies on the emergence of correlated swimming and the associated active turbulence, rather than solely on the vorticity of the condensate (Fig. 3). Instead, the expulsion correlates with the outward advective flow generated by correlated bacterial motion down the local cell-density gradient (Figs. 1, 2, and 4). Such alignment is consistent with the decrease in correlated bacterial motion at lower cell densities (28).

In addition to the outward force induced by correlated swimming in thick bacterial suspensions (h > h_0_), active turbulence can limit condensation through two additional mechanisms. First, by promoting advection of chemoeffectors (here, protons) it may limit the build-up of steep chemoeffector gradients. The plausibility of this effect can be estimated by comparing the timescale corresponding to the chemoeffectors diffusion across the condensate with the jet-like motion described in Fig. 2A, while taking into account the much larger diffusion coefficient of the molecular chemoeffector, relative to that of the beads (Fig. 2B). This analysis suggests that, although advection does not dominate over diffusion, it can nevertheless be significant. Second, correlations in the swimming direction of nearby cells can affect their run-and-tumble behavior. This effect has been suggested to either improve or impair chemotaxis in shallow gradients, depending on cell density (30). However, the bacterial fractional volume in the condensates (∼0.03 for h > h_0_) is far smaller than the densities that triggered reduced chemotaxis efficiency in ref. (30). Yet, this effect may partly account for the lower condensation efficiency observed in thin bacterial layers (h < h_0_) compared with porous environments (Figs. 3C and S8). On the other hand, correlations in bacterial swimming have also been suggested to enhance bacterial accumulation near local attractant source (31), which may be more relevant to bacterial condensates but would only enhance the condensation rather than broaden it.

In contrast to the consistent clockwise rotation observed here in condensates (Figs. 1–2), uniform bacterial suspensions confined in ∼30-µm deep and 45-µm wide wells, also exhibits vortex-like motion but with a random direction, whereas in wider wells the motion becomes chaotic (51). The clockwise vortex observed here in condensates arises from the basic rightward swimming tendency of the cells with respect to the radial velocity (Fig. 4C). This tendency is consistent with the circular motion exhibited by non-interacting individual cells swimming very close to glass surfaces (< 1 µM; Fig. S9) (52). However, bacterial condensates extend up to ∼60 µm from the surface and are far denser. Still, the asymmetry of the condensate with respect to the glass and the dense-hydrogel interfaces may break the left-right symmetry and promote clockwise rotation.

Finally, in contrast to the suppression of correlated swimming in porous hydrogel environments, correlated swimming was maintained in viscoelastic liquids containing mucin—albeit with larger vortex sizes—or in other synthetic analogues (38, 53). Interestingly, whereas mucin-containing media at neutral pH more closely resemble viscoelastic liquids, the agarose–hydrogel used here more closely resembles mucin media at acidic pH (54), which may be more relevant to gut physiological conditions. We suggest that the enhanced condensation observed in porous environments may facilitate the formation of more structured biofilm communities.

## Materials and Methods

### Strains and plasmids

We used *E. coli* MG1655 (VF6 (35, 55)) and its derivatives harboring deletion of *cheY* and *cheZ* (MK3) or deletions of all chemoreceptors and a dysfunctional *cheZ* gene (non-chemotactic run-and-tumble cells (AVE4 (35)). Cells expressed also a fluorescent mGFP (75 μM IPTG) or mCherry (5 mM IPTG) from a pTrc99a vector (pSA11).

### Cell growth and motility medium

Cells were grown over night in Tryptone Broth (tryptone extract 10 g/l and 5 g/l NaCl) supplemented with Ampicillin, diluted 100-fold into fresh medium with IPTG, and allowed to grow at 33.5 °C with shaking. Cells were harvested at OD_600_ 0.5 by centrifugation, gently washed twice in motility medium, and resuspended in motility medium to a final cell concentration of OD_600_ 0.4 ≈ 0.2·10^6^ cells/mm^3^ (unless otherwise mentioned). The motility medium (0.1 mM potassium phosphate, 0.94 mM lactic acid, 85.5 mM sodium chloride, 0.1 mM EDTA, and 1 μM methionine) was titrated to the desired pH with NaOH. This medium supports long-term bacterial motility but not cell proliferation. Experiments in porous medium (Fig. 3) were done gently mixing the cell suspension with agarose hydrogel (0.08–0.12% w/v) at 37 °C.

### The condensation setup

The setup is schematically illustrated in Fig. 1B (for more details see ref. (14)) A 3 % w/v agarose gel in motility medium was cast in a grade-2 titanium cylinder (6 mm high, 10.7 mm diameter) and sealed with a glass coverslip below and a custom top mold.

Molds were fabricated using a resin printer (Elegoo Mars 3) and had a thin cylindrical protrusion (radius of 5 mm in liquid and 3 mm in porous media; height ∼120 μm unless otherwise stated), which formed the bacterial layers. After the agarose solidified, the mold and the bottom cover were removed, and the agarose was dialyzed overnight in motility buffer and a second time, for an additional 4 h, in a fresh motility buffer just before the experiment (final dialysis ratio of 10^4^:1).

Cells suspensions (in liquid or dilute agarose) were then introduced into the shallow well and the setup was covered from both sides using a BSA-treated glass coverslips (1% bovine serum albumin in double-distilled water for 30 minutes) and secured in position. Experiments were performed at room temperature (24 °C).

### Fluorescence microscopy

The bacterial distribution was followed by using a Nikon Ti microscope equipped with a 20× (0.5 NA), 10× (0.45 NA), or 4× (0.2 NA) objective, a decoded controlled stage and a ‘perfect-focus’ system. Large images were obtained by stitching smaller images (approximately 2mm × 2mm each) with 20% overlap using a built-in function in NIS Elements software (Nikon). The bacterial distribution inside the well was extracted from the images by removing the measured background intensity, normalizing to the average intensity within each well, and multiplying by the initial (uniform) cell density. This results in the intensity being averaged over the vertical direction (z) (h): 𝜌 = 〈𝜌_3𝐷_〉_𝑧_. Radial averaging was done by measuring the fluorescence intensity around the center of focal points.

### Cell density estimates

Calibration of OD_600_ to cells/mm^3^ was done by plating serial dilution of cells after growth (OD_600_ 0.5), yielding OD_600_ 1 = (0.5±0.1)·10^6^ cfu/mm^3^ (N = 4). The total number of cells in a condensate (Fig. 4A) was calculated by integrating the density profile up to a 90% drop of the peak density. Calibration between OD_600_ to fractional volume was done by centrifuging a dense cell suspension and measuring the volume of the pellet; the dense pellet corresponded to approximately OD_600_ 500.

### Non-uniformity estimates

To determine the non-uniformity of the cell distribution within condensates from large still images (5 mm × 5mm), where temporal averaging is unavailable, a smooth radially-averaged version of each image was subtracted from the original ones. A high-pass filter (0-140 μm) was then applied to remove residual, large-scale intensity fluctuations.

We verified that the filter yields the same result as subtracting the time-averaged image in short time-lapse stacks (∼30 s) acquired in a smaller field of view (2 mm × 2 mm). Once filtered, non-uniformity was measured as the difference between the standard deviation of the intensity within a disk around the condensate center (600 μm diameter) and away from it.

### Fluorescent spheres

For PIV and particle tracking we used green fluorescent spheres (beads) with a 1μm diameter (FluoSpheres, ThermoFisher Scientific). Before use, spheres were thoroughly washed in motility medium to which Tween-20 (0.01%) was added. The spheres were then added to a cell suspension, which also contained Tween-20 (0.01%).

### Particle image velocimetry (PIV)

To capture velocity fields, time-lapse fluorescent images (2-20 frames per second) were captured and analyzed by PIV using PIVLab (56). To analyze the collective motion of cells, a filtering procedure was first applied to remove the large-scale bacterial condensation (see Non-uniformity estimates) and a nearest-neighbors smoothing scheme was applied to remove residual pixel-sized noise. Similarly-filtered results were obtained by subtracting the mean image from the stack. PIV analysis proceeded using multipass FTT window deformation. To avoid false detection, only vectors with a correlation coefficient larger than 0.5 were kept. The size of the interrogation windows was chosen to balance between the resolution of the extracted velocity field and the signal-to-noise ratio; changing them did not modify the results significantly. To analyze the motion of the fluorescent beads with a 4× objective, an initial mask was applied to ensure only particles within the focal plane were analyzed. This was possible since fluorescent beads within the focal plane seem smaller and brighter than beads in other planes. Similar results were obtained with a 10× objective with smaller depth of field (open markers in Fig. 3A and Movie S2). Before analysis, the median image was subtracted from the time lapse and thus only non-adhered beads were analyzed.

### Particle tracking and detection of jets

Time lapse images (2 frames per second, 4× objective) of the fluorescent beads were masked to ensure only particles within the focal plane are analyzed (see Particle image velocimetry (PIV)). Tracking was then performed using the TrackMate plugin for ImageJ. Detected tracks were exported and analyzed separately. Each track was smoothed using a Savgol-Golay filter (5 frames window) and velocities were estimated numerically. Jets were detected as temporal peaks in the velocity magnitude with a mean prominence of more than 0.5 μm/s. The length of each track (L) and it’s mean velocity were captured. If neighboring beads, less than L apart, exhibited a jet in the same direction (Δθ < 0.15), their jets were considered as correlated; the velocity of such jets were averaged over and considered as one. Due to the low density of beads in these experiments, such events were rare (7 of the 2362 detected jets).

### Expulsion of non-chemotactic bacteria

Experiments were performed using mixtures of chemotactic (red labeled) and non-chemotactic (NC, green labeled) bacteria mixed at a ratio of 100:1. Due to the low density of the NC cells, images were captured with a 10× and 20x objectives and density estimates were based on particle counting. A time series was captured, and the median image was subtracted from the sequence; thus, only moving cells were counted.

### Numerical solution of the model

Solutions and parameters were based on (14) and full details can be found in the Supp. Text. Briefly, the dynamics of cells was modeled in 2D assuming radial symmetry and the dynamics of protons was modeled in 3D assuming cylindrical symmetry. Proton uptake in the bacterial layer was accounted for by imposing absorbing boundary conditions for the protons at the bacterial layer interface. Numerical solutions were obtained by calculating spatial derivatives with 2^nd^ order accuracy and using a 4^th^ order Runge-Kutta method for time integration.

## Author Contributions

NL and AV designed the experiments; NL performed the experiments and theoretical calculations; NL and AV analyzed the data and wrote the manuscript.

## Acknowledgments

We thank Alexander Feigel (Hebrew University), Igor Aronson (Penn. State University) and Gabriel Amselem (École Polytechnique) for helpful conversations. We thank the Milner Foundation for partly funding NL’s fellowship. This work was supported by the Israeli Foundation of Sciences and Humanities and by a research grant from the U.S.-Israel Binational Science Foundation.

## Data Availability Statement

All data is available upon request.

