## Supplementary material for "Collective bacterial condensation is fundamentally constrained by the emergence of active turbulence": SI Appendix

**Supplementary Information**

SI Figures (pages 2-7)

SI Model (pages 8-13)

SI Movies (page 14)

### SI Figures

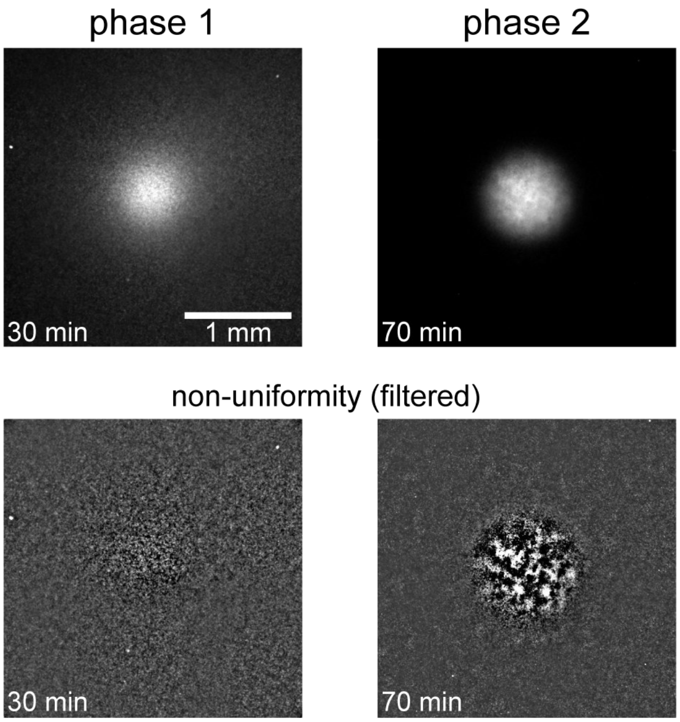


**Figure S1. Condensates are smooth during Phase 1.** Condensation experiments were performed in a liquid environment (data from Fig. 1). Shown are snapshots of a representative condensate during the final stages of Phase 1 (left) and in Phase 2 (right). Top: original fluorescence data displayed with adjusted grayscale ranges for clarity (left: 0–1500; right: 0–7000). The peak density was $1.44\times10^6$ cells/mm$^3$ in Phase 1 and $6.69\times10^6$ cells/mm$^3$ in Phase 2. Bottom: corresponding images after non-uniformity filtering (see Materials and Methods).

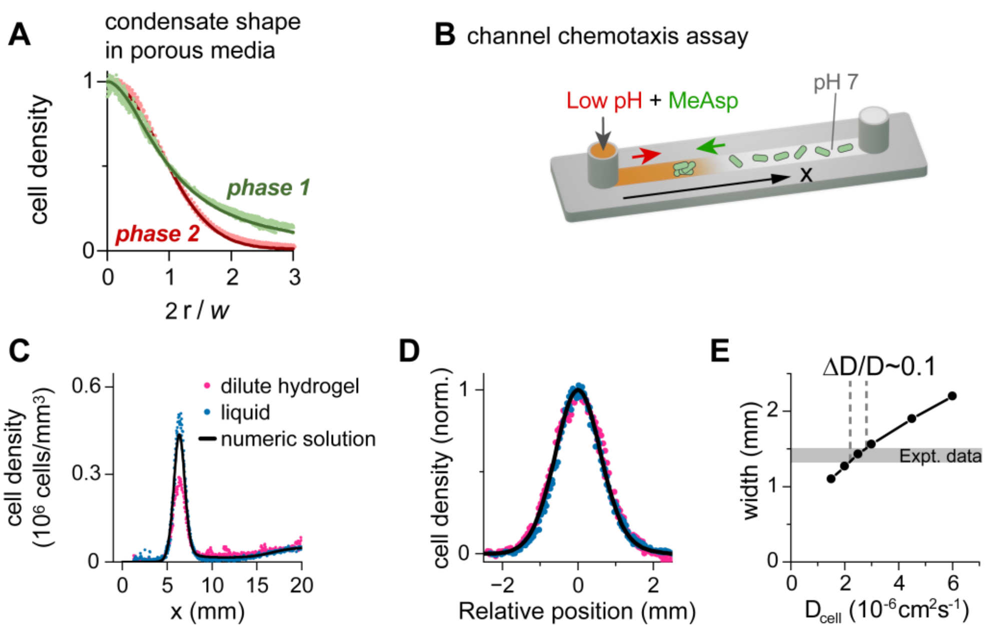


**Figure S2. The inherent chemotactic performance of E. coli is similar in liquid and in dilute hydrogel. (A)** Condensate shape in hydrogel during phases 1 and 2. Condensation experiments were performed in a dilute hydrogel environment (Fig. 3). The radially-averaged cell density profiles of condensates in Phase 1 and Phase 2 are plotted. Density profiles were normalized by subtracting the baseline cell density and dividing by the peak density; radial coordinates were normalized by the condensate half-width ($w/2$). Notably, during Phase 1 the condensate shape in hydrogel (light green markers) is similar to the corresponding shape in liquid (green line), showing that the cells' chemotactic performance is similar. **(B-E)** Chemotactic performance at low cell density. **(B)** Schematic of the channel setup. Following ref. (21), cells were uniformly distributed in a long channel and a source containing α-methyl-aspartate (MeAsp; attractant) adjusted to pH 4.5 (repellent) was introduced to one end. Chemoeffectors were allowed to diffuse and the cell density was measured by fluorescence microscopy at various times. To avoid collective condensation 1mM potassium was added to the motility medium **(C)** Cell density profiles at t = 4h in liquid and in dilute agarose gel (0.08%) are shown for two biological replicates each. Also shown is the numerical solution of the 1D chemotaxis model in response to a constant source of conflicting effectors (21):

$$\frac{\partial\rho}{\partial t} = D\frac{\partial^2\rho}{\partial x^2} - \frac{\partial}{\partial x}\left(\rho\frac{\partial}{\partial x}\left(\sum_{i=pH,MeAsp} \alpha_i \cdot \ln\left(\left(k_{off}^i + c_i\right)/\left(k_{on}^i + c_i\right)\right)\right)\right)$$

The simulation parameters used are listed below. **(D)** Normalized cell density near the peak. The density was normalized by subtracting the baseline and dividing by the peak value, and plotted as a function of distance from the peak center. This normalization is required to compare gel and liquid data, since the gel traps a fraction of cells, affecting the peak height and baseline but not the accumulation profile of motile cells. Notably, data in liquid and gel collapse together with the model onto a single curve. **(E)** Effect of cell diffusivity on accumulation width. To estimate the impact of enhanced active diffusion in liquid, the chemotaxis model was solved while varying only the cell diffusion coefficient. The resulting widths were compared to the experimental range (gray shaded). Together, these experiments show that the ratio between chemotactic bias and active diffusion is similar in the two environments. Simulation parameters:
$D = 3 \cdot 10^{-6}\ cm^2 s^{-1}$ (adapted from (14) to 1D), $\alpha_{pH} = 30 \cdot 10^{-6}\ cm^2 s^{-1}$ , $k_{off} = 10^{-3} M\ (pH\ 3), k_{on} = 10^{-6}\ (pH\ 6)$ ((14) as in Tabel S1.), and $\alpha_{MeAsp} = 25 \cdot 10^{-6}\ cm^2 s^{-1}$ , $k_{off} = 20\mu M, k_{on} = 20 mM$ (21). $c_i(x,t)$ was estimated by $c_i(x,t) = c_i^0 \cdot erfc(x/\sqrt{4D_i t})$ with $c_{pH}^0 = 10^{-4.5} M\ (pH\ 4.5)$, $D_{pH} = 2.5 \cdot 10^{-6}\ cm^2 s^{-1}$ and $c_{MeAsp}^0 = 10\ mM$, $D_{MeAsp} = 7.5 \cdot 10^{-6}\ cm^2 s^{-1}$ (21).

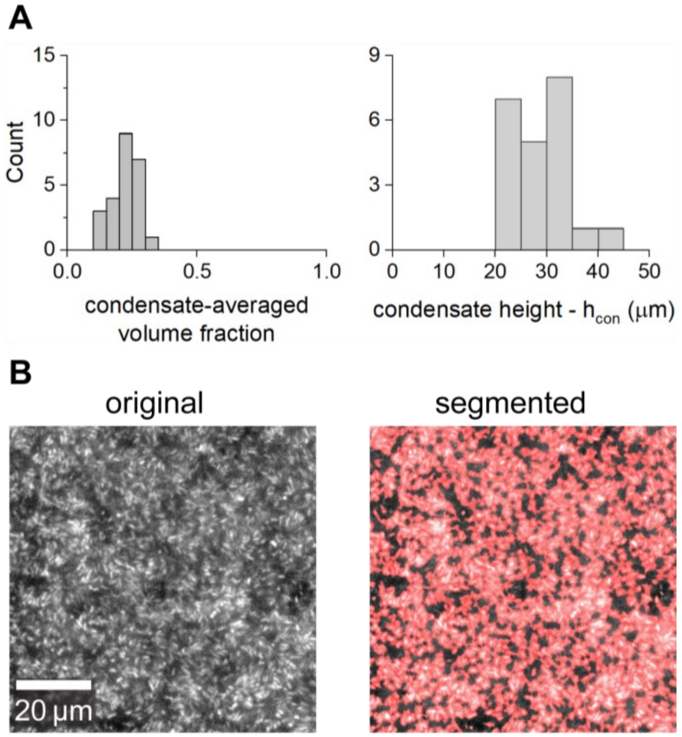


**Figure S3. Fractional volume in porous-media-embedded condensates.** Condensation experiments were performed in a dilute agarose environment (0.08-0.12% w/v; data from Fig. 3) and the fractional volume was estimated in two ways: **(A)** By calibration between the fluorescence intensity to fractional volume (see Cell density estimates in Materials and Methods) and **(B)** By measuring the covered area near the glass coverslip. **(A)** Histogram of the intensity-estimated condensate-averaged volume fraction (left plot; avg. ± SD = 0.23 ± 0.05, N = 24). The density with the condensate half-width was averaged and also corrected for relative occupation of the condensate within the well: cell density was measured by integration over the z-coordinate and corrected for the fact that condensates sedimented to the bottom 20-30 μm of the well (right plot; $h_0^{gel}$ = 26±6 μm, N = 22). The peak volume fraction was approximately 25% higher. **(B)** Area fractional at the condensate peak near the bottom surface. Fluorescence image at the condensate peak was captured using a 100x objective lens near the glass (bottom) surface. Left – original image; Right – segmented image showing the cell-covered region in red. In this experiment, the covered area at the peak was ~0.5-0.7. Note that the initial (uniform) fractional volume was ~$10^{-3}$.

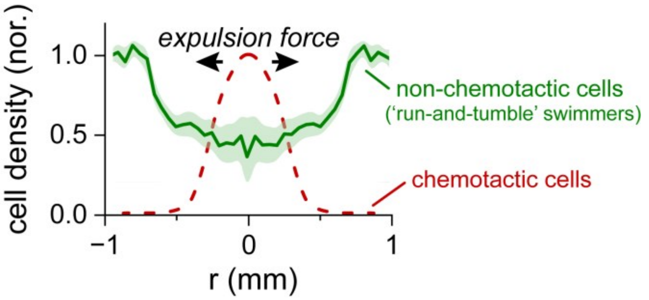


**Figure S4. Expulsion of non-chemotactic, run-and-tumble swimmers by condensing bacteria.** Experiments were performed as in Fig. 3D, but with non-chemotactic *E. coli* that exhibit run-and-tumble motility rather than smooth-swimming (strain AVE4 (35)). These cells lack all chemoreceptors and carry a CheZ mutation that induces random tumbling events. Green line: mean of 3 independent experiments, green shade: SD around the mean. The density of the chemotactic cells was normalized relative to the peak cell density; non-chemotactic cells were normalized relative to the density away from the peak. Expulsion in this strain is less pronounced than in the smooth-swimming cells (Fig. 3D), likely because these run-and-tumble cells exhibited weaker tendency to accumulate at the lower region of the bacterial layer, making them less exposed to forces in the condensate plane.

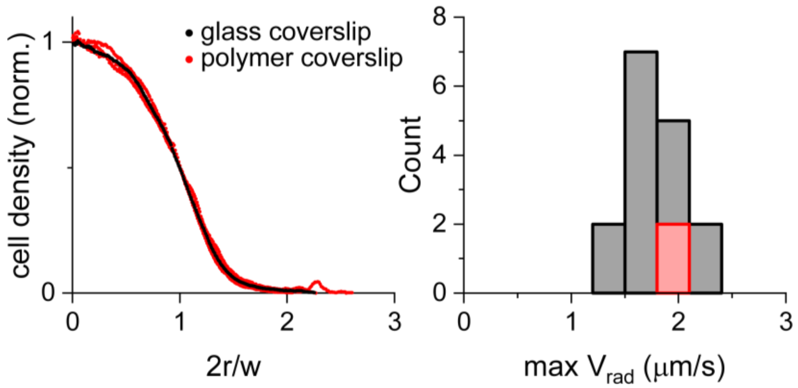


**Figure S5. Condensation experiments with oxygen permeable coverslips.** Condensation experiments were performed as in Figs. 1-2, except that oxygen-permeable ibidi polymer coverslips (cat. no. 10814) were used (pre-treated with 1% BSA as usual). Results in experiments with the oxygen-permeable coverslips (red, N=3) are contrasted with ones using glass coverslips (black). Left – normalized cell density profiles during Phase 2 (normalized as in Fig. 1D: peak = 1, baseline = 0 and r was divided by the half-width, w/2). Right – histogram of the maximal radial velocity in several experiments (black – data from Fig. 3A with $h > h_0 = 60$ μm). The initial cell density in these experiments was $0.2 \cdot 10^6$ cells/mm$^3$ (OD 0.4), below the threshold for ring formation. Evidently, the polymer coverslips did not significantly influence condensation. These results agree with previous results showing that cells lacking the Aer receptor condensed like the wildtype strain in similar experiments (14).

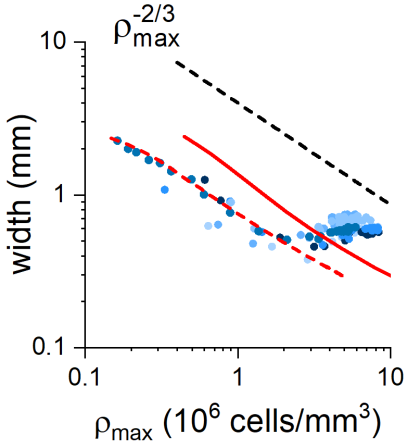


**Figure S6. The basic chemotaxis model captures the power-law between $\rho_{max}$ and $w$.** The basic chemotaxis model (Eqs. 1-2) was solved without the additional cell-induced advection term (Eq. 3). Once a condensate started forming $\rho_{max}$ and $w$ displayed a power law relation $w \sim (\rho_{max} - \rho_0)^{-M}$ with $M \approx 0.66 - 0.7$ as in the experiments (Fig. 1D). Note that using the parameters from Table S1 (SI Model) resulted in a pre-factor slightly different from the experiments (solid red line, shifted relative to the data). The slope of the of the power law seems robust in our model. Specifically, $\pm 50\%$ changes in the proton removal rate ($\gamma$), the initial cell density ($\rho_0$), the bacterial layer thickness ($h$), or the lateral dimension of the system did not change the power law but only its pre-factor. Using a smaller lateral system size (d = 3mm) resulted in the dashed red line. This discrepancy in the pre-factor may stem from the fact that unlike the model, in the experiments, condensation is initiated through cell-density fluctuations rather than boundary conditions alone (14).

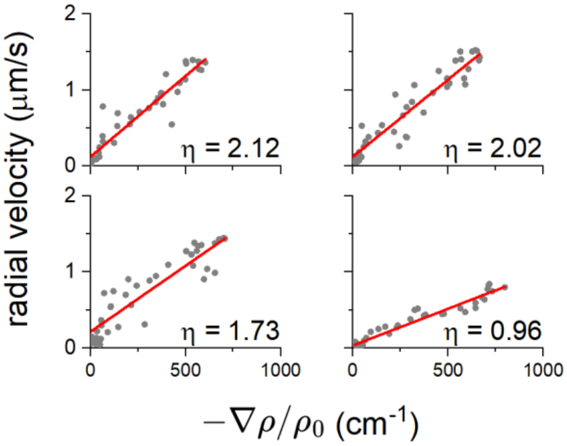


**Fig. S7. Radial fluid velocity vs cell density gradient.** The radial fluid velocity $(v_{rad})$ is plotted against minus the cell density gradient $(-\partial\rho/\partial r)$ divided by the initial cell density $\rho_0$ = $OD_{600}$ 0.4 (~ $0.2\cdot10^6$ cells·mm$^{-3}$). Red lines show linear fits of the form $v_{rad} = -\eta\nabla\rho/\rho_0$ with η in units of $10^{-7}$ cm$^2$·s$^{-1}$, providing $\eta = 1.7 \pm 0.5 \cdot 10^{-7}$ cm$^2$·s$^{-1}$. Variability between fits is most likely due to the low SNR of $\rho_0$ in the red channel in these experiments, which stretches the horizontal axis by a constant. Data was taken from Fig. 2D and presented here in physical units.

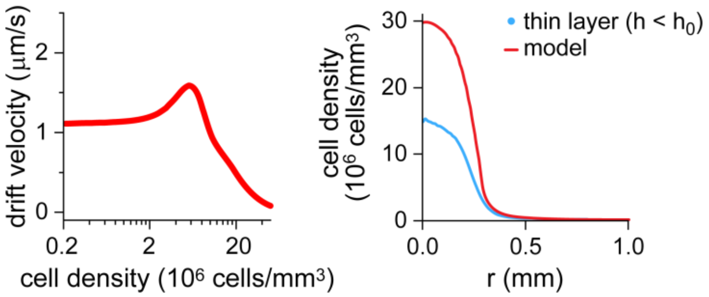


**Figure S8. Chemotaxis deficiency due to active turbulence may limit cell condensation at high volume fractions.** Left - data from Fig. 3a in ref. (30) was used to estimate the density-dependence of the chemotaxis drift velocity. Fractional volume $(\phi)$ was converted to cell density by setting ϕ=1 ↔ $250\cdot10^6$ cells/mm$^3$ (see Materials and Methods) As noted in (30), the chemotaxis drift had a slight increase at intermediate cell density. Fitted function: $v_0 \cdot f(\rho)$ with $v_0 = 1.1\ \mu m\ s^{-1}$ and $f(\rho)\left(1-\frac{\rho^{2.5}}{\rho^{2.5}+91^{2.5}}+0.5\cdot\exp\left(\frac{(\rho-30)^2}{236}\right)\right)$. Right – the basic chemotaxis model was solved assuming that the chemotaxis coefficient $\beta$ has the density dependence: $\beta(\rho) = \beta_0 f(\rho_{max})$. Note that by using the condensate peak density, $\rho_{max}$, we estimate here the maximal possible contribution of the effect. Red: numerical solution, light blue: data from Fig. 3 in thin bacterial layers. Here, the simulated initial density and the layer thickness were set to match the experimental conditions (h = 40μm, $\rho_0$ = $OD_{600}$ 0.8).

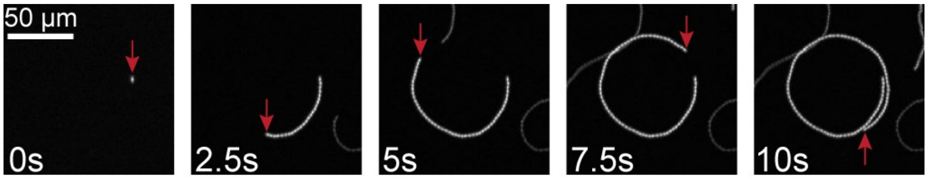


**Figure S9. Bacteria swim in circles near solid surfaces.** Smooth-swimming cells were introduced into the thin-layer assay (pH 7, 10 mM potassium phosphate), and their motion was captured using fluorescence time-lapse imaging. Shown are maximum-intensity projections of increasing duration (as labeled) for a representative cell. Red arrows mark the position of the cell. Consistent with previous observations (52), cells swimming near the bottom glass surface move in clockwise circles when viewed from beneath.

### 2. SI Model

This section contains the following subsections:

2.1 **The basic model.** Description of the basic chemotaxis model, without high-density corrections.

2.2 **Effects of collective swimming on bacterial condensation.** Description of the effects considered due to correlated swimming (radial outflows and chemotaxis deficiency).

2.3 **Phase 2 (quasi-steady state) solution.** Analytical solution to the shape of mature condensates (Phase 2). Related to Fig. 5A

2.4 **Expulsion of non-chemotactic cells**. Analytical solution for the distribution of non-chemotactic cells due to repulsion from condensates.

2.5 **Numerical scheme**.

#### 2.1 The basic model

Following previous work (14, 40), the basic model used here is based on the Keller-Segel equations (39):

$$\begin{cases} \dfrac{\partial \rho}{\partial t} = \vec{\nabla} \cdot \left( D \vec{\nabla} \rho - \rho \vec{v}_D \right) & (1) \\ \dfrac{\partial c}{\partial t} = D_c \nabla^2 c - r_H \rho c & (2) \\ \vec{v}_D = \beta \vec{\nabla} f \quad ; \quad f = \ln\left( \dfrac{c + K_{off}}{c + K_{on}} \right) & SE.1 \end{cases}$$

The dynamics of the cells ($\rho$) and protons ($c$) are characterized by Eqs. (1) and (2), respectively. In our model, the number of bacteria is conserved, and bacteria move due to random motion, characterized by an active diffusion coefficient $D$, and due to chemotaxis-induced drift ($\vec{v}_D$). Protons also diffuse, with a diffusion coefficient $D_c$, but their number is not conserved; protons are consumed by the bacteria.

Following ref. (41) the chemotaxis drift velocity is described by Eq. SE. 1, where $K_{on}$ and $K_{off}$ are the effective dissociation constants for the receptors in the ON or OFF states, and $\beta$ is the chemotaxis coefficient. Note that this term expresses the log-sensitivity of cells within the dynamic range of $K_{off} \ll c \ll K_{on}$. Finally, $r_H$, which determines the proton uptake, was set to $r_H = \gamma c^{1/2}$ as observed in experiments (14).

In our setup cells are confined to a thin (120 μm) layer, with typical diffusion time across the layer of about ~20 seconds. Thus, the equation for the cell density (Eq. (1)) was solved in two dimensions assuming radial symmetry. In contrast, protons can diffuse freely throughout the chamber, and thus their dynamics (Eq. (2)) was considered in three dimensions assuming cylindrical symmetry.

Proton consumption by the bacteria was modeled by imposing absorbing boundary conditions at the bacterial layer interface, with a proton flux $\overrightarrow{J_c}$ given by the 2D cell density and the bacterial proton consumption rate $r_H$:

$$\vec{J_c} \cdot \hat{z}\left(z = 0, r \leq \frac{d}{2}\right) = h \cdot r_H \cdot \rho \qquad SE.2$$

where $h$ is the thickness of the bacterial layer, set to 120 μm, and $d$ is the well diameter. All other boundary conditions were set to ensure no cell or proton flux at their respective boundaries. The initial conditions were set such that bacteria and protons were uniformly distributed (cells at $OD_{600}$ 0.4 ≈ $0.2 \cdot 10^6$ cells/$mm^3$ within the bacterial layer and protons throughout the chamber at pH 5.2).

Notably, in this model, condensation is only limited by the sensory dynamic range; cells condense until the proton concentration at the peak reaches $c \sim K_{on}$. The resulting dynamics appear to be consistent with experiments where no collective motion is observed but the model does not capture the data at high cell counts (40 and Fig. 5). In addition, the peak density and the width of calculated condensates seem more sensitive to the lateral system size and the initial cell density than the experimental condensates, further suggesting that the basic model lacks a limiting mechanism at high cell counts (40). These effects are summarized in Fig. 5D.

### 2.2 Effects of collective swimming on bacterial condensation

To study the possible implications of collective swimming on bacterial condensation, we made modifications to the basic model. Aiming mostly to show that the proposed mechanisms are relevant in terms of their potential effect, our modeling remains as simple as possible. The effects of cell-induced advective flow and of chemotaxis deficiency due to collective motion were tested separately.

**Cell-induced fluid motion.** To test whether the cell-induced radial convective fluid flow (Fig. 2) can limit condensation, we modified the drift velocity (Eq. SE. 1):

$$\vec{v}_D = \beta\vec{\nabla}f + \vec{U} \qquad SE.3$$

Phenomenologically, we found that $\vec{U}$ was consistently proportional to the gradient of the cell density (Figs. 2E, 4C and S7), and was therefore approximated by:

$$\vec{U} = -\eta\vec{\nabla}(\rho/\rho_0) \qquad SE.4$$

The data in Fig. S7 was fitted with $\eta = 1.7 \pm 0.5 \cdot 10^{-7}\ cm^2 s^{-1}$ and $\rho_0$ = $OD_{600}$ 0.4 ≈ $0.2 \cdot 10^6$ cells/mm$^3$. To account for the abrupt transition between Phase 1 and Phase 2 we introduced a sigmoidal function of the maximal cell density such that
$\vec{U} = -\sigma(\rho)\eta\vec{\nabla}(\rho/\rho_0)$ using the following function:

$$\sigma(\rho) = \frac{1}{1 + \exp\left(-\frac{k}{\rho_0}\left(\rho_{max} - \rho_{\frac{1}{2}}\right)\right)} \qquad SE.5$$

where $k = 3$ and $\rho_{1/2}$ = $25\rho_0 \approx 5 \cdot 10^6$ cells·mm$^{-3}$. See below the analytical solution of the modified model in the quasi-steady state (section 2.3).

**Chemotaxis deficiency.** Collective swimming was previously suggested to modulate the chemotaxis efficiency of the bacteria (30). To model this effect, we added a density-dependence to the chemotaxis efficiency of the bacteria, i.e., $\beta = \beta(\rho_{max})$, where $\rho_{max}$ is the peak cell density. We estimated this cell density dependence of $\beta$ based on (30) and incorporated it into the model (Fig. S8). Note that by using $\rho_{max}$ we evaluate the maximal contribution of this effect.

**2.3 Analytical solution for the condensate shape with cell-induced radial flow (Phase 2)**

We add Eqs. SE.4 and SE. 5 to the basic model to obtain:

$$\frac{\partial\rho}{\partial t} = \vec{\nabla}\cdot\left(D\vec{\nabla}\rho - \rho\big(\vec{v}_D - \sigma(\rho)\eta\vec{\nabla}(\rho/\rho_0)\big)\right) \qquad SE.6$$

At the quasi-steady state, we can integrate equation SE. 6 once:

$$D\vec{\nabla}\ln(\rho) + \eta\sigma(\rho)\vec{\nabla}(\rho/\rho_0) = -\beta\vec{\nabla}\ln(c+k) + C_1 \qquad SE.7$$

$C_1$ is an integration constant. Since all gradients vanish far from the condensate we set $C_1 = 0$. For $\rho_{max} \gg \rho_{1/2}$ at the steady state we find $\sigma = 1$ which simplifies the solution. Otherwise, except for special cases exactly at the transition, we have $\sigma = 0$ and we recover the known solution without any fluid drift (40). Using $\sigma = 1$ and integrating once again leads to:

$$Dln\rho + \eta(\rho/\rho_0) = -\beta\ln(c+k) + C_2 \qquad SE.8$$

Which simplifies to

$$\frac{\eta\rho}{\rho_0 D}exp\left(\frac{\eta\rho}{D}\right) = \frac{\eta\rho^*}{\rho_0 D}\left(\frac{c+k}{c(r=0)+k}\right)^{-\beta/D} \qquad SE.9$$

We recognize that the equation can be inverted using the W function:

$$\rho = \frac{D}{\eta}W\left(\frac{\eta\rho^*}{D}\left(\frac{c/k+1}{c(r=0)/k+1)}\right)^{-\frac{\beta}{D}}\right) \qquad SE.10$$

The functional shape of $c(r)$ can be approximated using the asymptotic assumption of $c(r\to 0)\sim c_0 + \alpha r^2$ and $c(r\to\infty) = c_\infty$ using the following ansatz (37):

$$c(r) = c_0 + (c_\infty - c_0)\tanh^2(r/\ell) \qquad SE.11$$

The parameters here must be determined by solving the equation for the protons (see (40)). Plugging SE.11 into SE.10 fits the experimental data, albeit with several free parameters.

#### 2.4 Analytical solution for the repulsion of non-chemotactic cells from condensates

Writing SE.6 for non-chemotactic (NC) cells in the presence of wildtype (WT) condensed cells, we find ($\sigma = 1,\ \beta = 0$):

$$\frac{\partial \rho_{NC}}{\partial t} = \vec{\nabla} \cdot \left(D_{NC}\vec{\nabla}\rho_{NC} + \rho_{NC}\eta\vec{\nabla}(\rho_{WT}/\rho_0)\right) \qquad SE.12$$

Where we've implicitly neglected the contribution of the NC cells to the radial outflows due to their low density. At the quasi-steady state, we can integrate the equation once:

$$D_{NC}\vec{\nabla}\rho_{NC} = -\rho_{NC}\eta\vec{\nabla}\left(\frac{\rho_{WT}}{\rho_0}\right) + C_1 \qquad SE.13$$

Since both gradients vanish far from the condensate center, we set $C_1 = 0$. Rearranging the equation we find:

$$\vec{\nabla}\ln(\rho_{NC}) = -\frac{\eta}{D_{NC}}\vec{\nabla}\left(\frac{\rho_{WT}}{\rho_0}\right) \qquad SE.14$$

Another integration leads to:

$$\rho_{NC} = \rho_{NC}^{\infty}\exp\left(-\frac{\eta}{D_{NC}}\frac{\rho_{WT}}{\rho_0}\right) \qquad SE.15$$

where the integration constant $\rho_{NC}^{\infty}$ is the density of NC cells far from the condensate center. In the main text, we normalize the density by setting $\rho_{NC}^{\infty} = 1$.

### 2.5 Numerical scheme

Eqs. (1) and (2) were solved numerically by calculating the spatial derivatives with an accuracy of $\mathcal{O}(\Delta x^2)$ and integrating over time using an explicit 4th order Runge-Kutta method with an accuracy of $\mathcal{O}(\Delta t^4)$. The scheme was tested as described before (14, 40). Briefly, we verified convergence by ensuring that the output remained unchanged when further reducing Δt and by comparing the numerical results in special cases where analytical solutions are known.

| Parameter | Notes |
|---|---|
| $D = 5 \cdot 10^{-6}\ cm^2 s^{-1}$ | Active cell diffusion coefficient |
| $\beta = 30 \cdot 10^{-6}\ cm^2 s^{-1}$ | Chemotaxis coefficient |
| $K_{on} = 10^{-6}\ M\ (pH\ 6)$ | Effective dissociation constant (ON state) |
| $K_{off} = 10^{-2}\ M\ (pH\ 2)$ | Effective dissociation constant (OFF state) |
| $D_c = 5 \cdot 10^{-6}\ cm^2 s^{-1}$ | Proton diffusion coefficient |
| $\gamma = 0.035\ mM^{-1/2} s^{-1}\ (OD\ 0.4)^{-1}$ | Proton uptake rate coefficient |
| $\eta = 1.7 \cdot 10^{-7}\ cm^2 s^{-1}$ | Coefficient of cell-induced flows |

**Table S1.** Summary of the parameters used in the numerical solutions of the model. The parameters were estimated based on available data; detailed calculations can be found in Table M1 in (14) and Fig. S7.

### SI Movies Videos

**Movie S1 – Collective swimming during bacterial condensation.** Experiments were performed as described in Fig. 1. The video shows a mature condensate (Phase 2) recorded using fast time-lapse imaging (5 frames per second). Left – original fluorescence data. Right – filtered data (see Non-uniformity estimates in Materials and Methods for filtering procedure).

**Movie S2 – Fluid flows in and above the condensate plane.** Experiment was performed as described in Fig. 2. The motion of green-fluorescent beads was recorded using fast time-lapse imaging (20 frames per second) at various heights (z coordinate) using a 10x objective (depth of field ~ 6 μm); The median image of the time-lapse at each height was subtracted from all frames in that height. Shown at x15 real speed. Notably, for $z < h_0$ beads flow outward from the condensate while for $z > h_0$ beads flow inward.

**Movie S3 – Collective motion persists in thin bacterial layers ($h < h_0$).** Experiment was conducted with a thin bacterial layer ($h < h_0$) as described in Fig. 3A. Time-lapse images were captured at 14 fps using a 10x objective. The median image of the movie was subtracted from each frame. Shown at x3 real time.

**Movie S4 – Motility of bacteria within a condensate embedded in dilute hydrogel.** Experiment was conducted in dilute agarose as described in Fig. 3B. The motion of bacteria was captured using time-lapse imaging (8 frames per second) at high magnification (100x) inside the condensate. The median image of the movie was subtracted from each frame.
